# Induced tolerance to UV stress drives survival heterogeneity in isogenic *E. coli* cell populations

**DOI:** 10.1101/2025.05.14.654146

**Authors:** Shunsuke Ichikawa, Midai Tanoue, Junto Takeuchi, Eri Matsuo, Yasuhito Shimada, Abhyudai Singh

**Affiliations:** Faculty of Education, Mie University, 1577 Kurimamachiya-cho Tsu, Mie, 514-8507, Japan; Zebrafish Research Center, Mie University, 1577 Kurimamachiya-cho Tsu, Mie, 514-8507, Japan; Department of Integrative Pharmacology, Graduate School of Medicine, Mie University, 2-174 Edobashi Tsu, Mie, 514-8507, Japan; Department of Electrical and Computer Engineering, Biomedical Engineering and Mathematical Sciences, Center of Bioinformatics and Computational Biology, University of Delaware, Newark, Delaware 19716, USA

**Keywords:** *Escherichia coli*, persister, primed cells, ultraviolet light irradiation, Modified Luria-Delbrück fluctuation test

## Abstract

The emergence of transiently tolerant bacterial subpopulations challenges our understanding of stress tolerance mechanisms. While much is known about antibiotic tolerance, it remains unclear whether similar mechanisms contribute to survival under ultraviolet (UV) stress. Here, we employed a modified Luria–Delbrück fluctuation test to investigate the presence of pre-existing UV-tolerant subpopulations in *Escherichia coli*. Our results showed no significant evidence of pre-stress UV tolerance. Instead, the data suggest that survival is primarily driven by inducible DNA repair responses activated after UV exposure. Furthermore, sequential low-dose UV exposures yielded higher-than-expected survival, suggesting that transient tolerance can be induced following initial UV exposure, likely through active DNA repair processes. These findings indicate that *E. coli* survives UV stress via an induced, rather than pre-existing, mechanism of tolerance.

## Introduction

The emergence of tolerant cells in clonal bacterial populations challenges the traditional view of cellular homogeneity and calls for a reassessment of how genetically identical cells respond to environmental stress. This phenomenon is not merely the result of random fluctuations; rather, it reflects a survival strategy that enables populations to thrive amid unpredictable environmental changes. Understanding the reversible, non-genetic processes that confer tolerance is therefore essential for advancing our knowledge of microbial ecology, evolution, and the development of strategies to combat antibiotic tolerance ^1–3^. A transiently tolerant state allows cells to switch between sensitive and tolerant phenotypes in response to specific environmental cues. This phenotypic plasticity represents a bet-hedging strategy in which a subpopulation pre-adapts to anticipated stress conditions without bearing the long-term costs of permanent genetic changes ^4–6^.

At the molecular level, the reversible acquisition of a tolerant phenotype is driven by stochastic fluctuations in gene expression and metabolic processes ^7,8^. These random variations, known as metabolic noise, allow isogenic cells to spontaneously adopt alternative functional states ^9,10^. Importantly, epigenetic mechanisms can help stabilize these transient states over several cell divisions, providing a measurable degree of cellular memory that extends beyond immediate environmental triggers ^11^. This interplay between stochasticity and epigenetic regulation allows microbial populations to maintain a flexible yet robust response to stress, thereby enhancing their long-term fitness in fluctuating environments ^12^.

To study reversible, non-genetic tolerance states, researchers have employed a modified version of the classical Luria-Delbrück fluctuation test (FT). Originally developed to estimate mutation rates, the FT showed that phage resistance in bacteria arises from spontaneous mutations rather than being induced by phage exposure. In the classical test, single bacterial cells are grown into clones and then exposed to phages. If mutations were induced by the virus, the number of resistant cells per clone would follow a Poisson distribution. Instead, Luria and Delbrück observed a highly variable, non-Poisson distribution, indicating that mutations occurred spontaneously before selection ^13^. While the original FT focused on irreversible genetic changes, modern adaptations have extended its use to investigate reversible phenotypic switching ^14^. It has been applied to detect transient, non-genetic cell states—such as reversible switches between drug-sensitive and drug-tolerant phenotypes— that exist even prior to stress exposure ^15^. Measuring survival variability among clones allows researchers to infer switching dynamics and rates ^16^. This approach enables quantitative detection of pre-stress tolerant subpopulations and provides insight into how non-genetic heterogeneity influences microbial stress responses.

Recent studies have shown that even in genetically homogeneous cancer cell populations, rare subpopulations can transiently express tolerance markers like EGFR and NGFR ^17,18^. Using fluctuation tests alongside single-cell techniques, researchers have revealed that such primed states can persist for several divisions, reflecting transcriptional memory ^19–22^. Similar approaches in viral systems have demonstrated that infection susceptibility is influenced by preexisting cellular states rather than random noise ^23,24^. These findings highlight the broader applicability of the fluctuation test in uncovering how non-genetic heterogeneity affects cell fate decisions.

The phenomenon of persister cells in bacteria represents a well-studied example of transient tolerance ^25–28^. These cells exemplify non-genetic heterogeneity and are believed to contribute to the failure of antibiotic therapies ^29^. Rahman et al. recently showed, via transcriptome profiling, that *Escherichia coli* persister cells challenged with ampicillin remain metabolically active—exhibiting widespread stress-responsive gene upregulation despite growth arrest—thereby challenging the long-held view that persistence equates to dormancy ^30^. The FT has revealed that shifts in trehalose metabolism—termed the trehalose catalytic shift—enhance this metabolic heterogeneity and increase the frequency of drug-tolerant persisters in *Mycobacterium tuberculosis* ^31^. In *Salmonella enterica*, subpopulations with heterogeneous flagellar expression exhibit significantly higher persistence to ciprofloxacin and streptomycin ^32^. Intracellular iron levels function as a heritable iron memory that modulates both swarming motility and antibiotic tolerance, predisposing cells to multidrug tolerance in *E. coli* ^33^. In addition, persisters have been shown to arise from stochastic drops in ATP levels, with low-energy subpopulations demonstrating increased survival under antibiotic treatment ^34^.

Primed cells, unlike persisters, continue to divide while maintaining transient tolerance for several generations. Hossain et al. have identified primed cells in *E. coli* clonal populations ^35^. Guha et al. have demonstrated that primed cells are not exclusive to *E. coli* but also exist in Gram-positive bacteria such as *Priestia megaterium* and *Bacillus subtilis*, where they contribute to increased antibiotic tolerance ^36^. Cross et al. reported that clinical isolates of *Klebsiella pneumoniae* harbor a small subpopulation with multi-generational tolerance to meropenem, suggesting that priming mechanisms are present even in clinically significant pathogens ^37^. Mathematical models examining the dynamics of primed cells suggest that they proliferate at rates close to non-primed cells, distinguishing them from slow-growing or dormant cells ^35^.

While primed cells have been well-characterized under antibiotic stress, their role in UV stress response remains unclear. Clonal populations of cells are known to exhibit variation, and from the perspective of DNA repair, this variation is likely also involved in UV stress tolerance ^38^. We previously reported the presence of rare *E. coli* cells that are temporarily tolerant to UV irradiation within clonal populations ^39^. Importantly, this UV stress tolerance does not the result from permanent genetic alterations ^40,41^; rather, it appears to be a stochastic and reversible state that enables a subset of cells to survive in UV-rich environments. The modified Luria-Delbrück fluctuation test serves as a powerful tool in this context, enabling analysis of the distribution of tolerant colonies and inferring the underlying mechanisms of transient tolerance. In this study, we applied a modified Luria-Delbrück fluctuation test to determine whether survival under UV stress in *E. coli* is driven by pre-existing primed cells (Fig. 1). If survival relies on a pre-stress priming mechanism, we would expect subpopulations with enhanced UV tolerance before exposure. In contrast, if survival is predominantly induced post-exposure, any elevated tolerance would only emerge after an initial UV dose. Our findings ultimately reveal that *E. coli* primarily relies on an induced mechanism, rather than pre-existing primed cells, to survive lethal UV stress. Additionally, sequential low-dose UV irradiations yielded higher-than-expected survival, underscoring the importance of induced UV stress tolerance in mitigating cumulative damage. These results reveal a distinct mechanism underlying transient stress tolerance in *E. coli*, highlighting the critical role of induced responses to UV stress and offering new insights into how bacterial populations adapt to diverse environmental challenges.

**Fig. 1.**
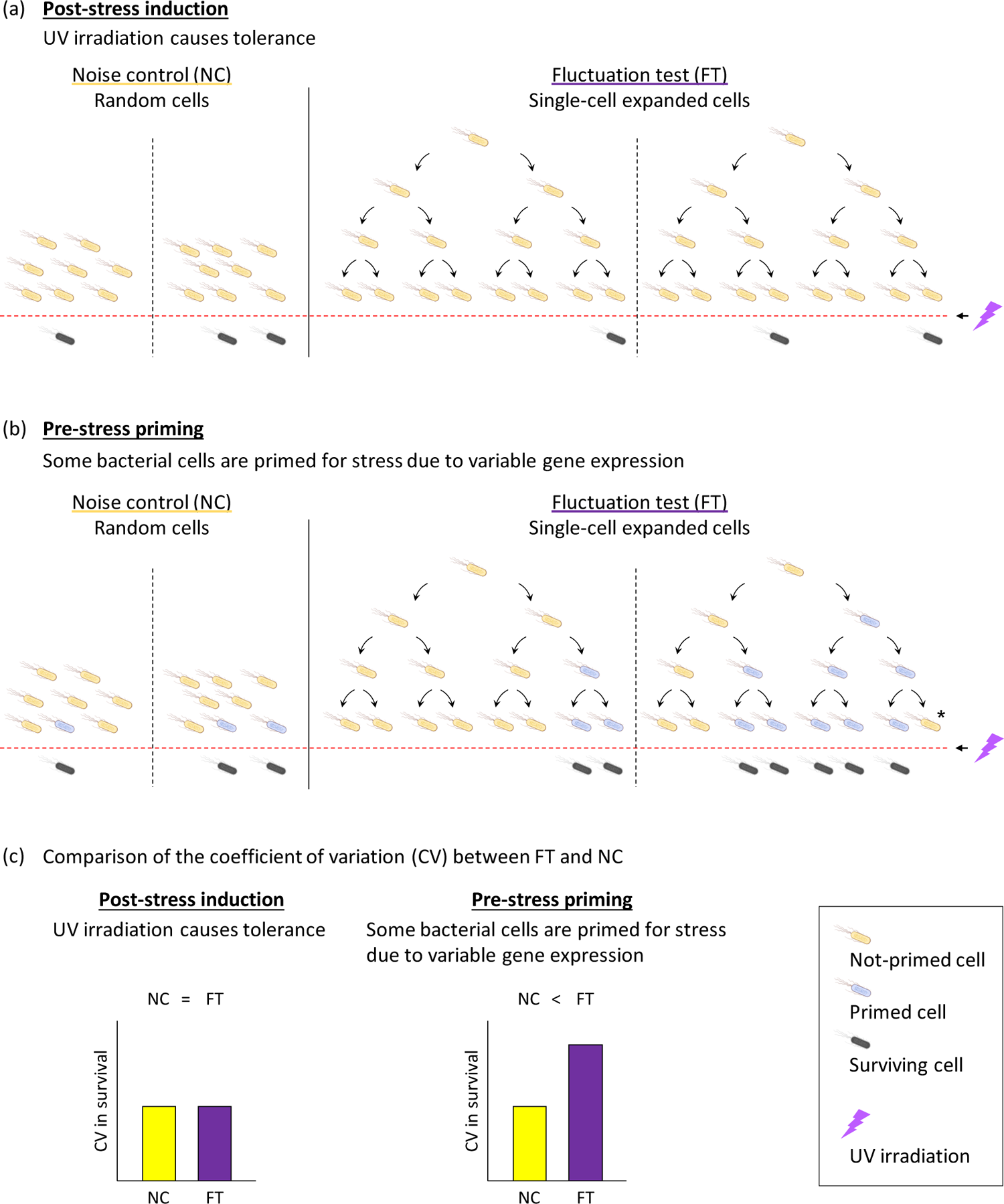
The modified Luria–Delbrück fluctuation test can reveal phenotypic variations among individual bacterial cells The modified Luria–Delbrück fluctuation test compares viabilities between single-cell expanded cell populations (FT) with random cell populations (NC) to detect variability arising from pre-existing tolerant cells. (a) If UV irradiation purely induces tolerant cells, their distribution across clones would follow a Poisson pattern. (b) Alternatively, if some cells are primed to respond to stress before UV irradiation, significant variations in the number of tolerant cells would be observed among clones. Primed cells exhibit reversible characteristics and may revert to a non-tolerant state after several generations, as indicated by asterisk. (c) If UV irradiation purely induces tolerant cells, no difference in tolerant cell variation is expected between FT and NC. However, if some cells are primed for stress tolerance before UV irradiation, greater variation in tolerance is observed among the clonal populations in FT compared to NC.

## Materials and methods

### Population-level variation of *E. coli* viability after UV irradiation (NC: noise control)

*E. coli* K-12 was first grown aerobically in Luria-Bertani (LB) broth at 37 °C. When cultures reached the logarithmic phase (OD_600_ of 0.3–0.5), they were diluted 1,000-fold in phosphate-buffered saline to yield 15 mL of suspensions (10^5^–10^6^ CFU/mL) in 55 mm diameter glass Petri dishes. A total of 12 samples were prepared for each experiment. A 262 nm UV-LED (Nikkiso Giken Co. Ltd., Ishikawa, Japan) was placed 32 mm above the suspension surface, which was stirred at 700 rpm during irradiation (Fig. 2).

**Fig. 2.**
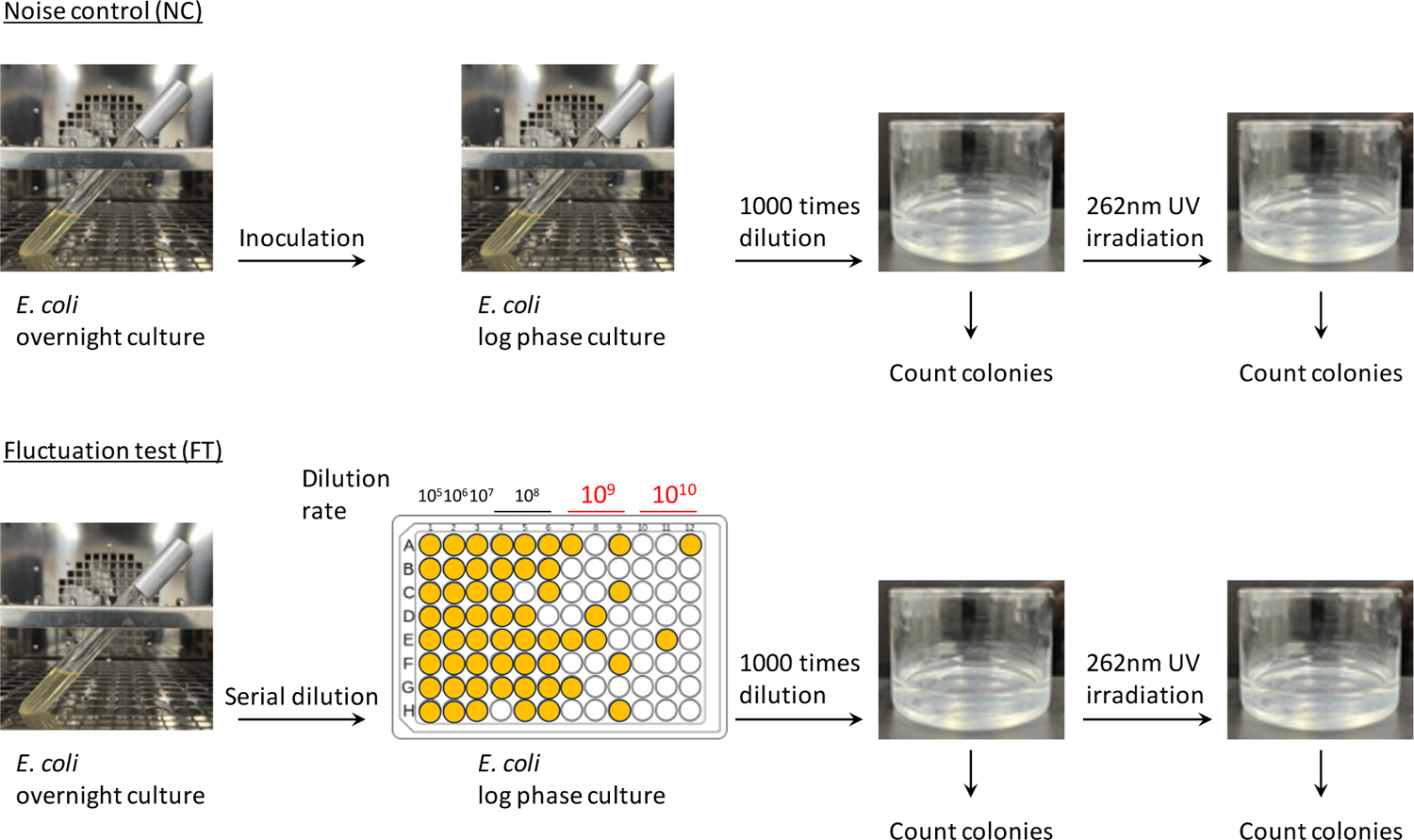
Experimental design for the fluctuation test by UV irradiation To examine tolerant cell variation in a single population, referred to as noise control (NC), *E. coli* cultures were grown to log phase, diluted into glass dishes, treated with UV, and plated before and after UV irradiation to determine viabilities. To examine tolerant cell variation in clonal populations originating from single cell level, referred to as FT.

After the UV irradiation, the treated cells were plated onto 3M Petrifilm Aerobic Count Plates (3M Company, MN, USA) and incubated at 35 °C for 48 hours. Viable cell counts were determined as colony-forming units (CFUs) per milliliter in the UV-treated samples (N). These counts were then normalized to those of untreated samples (N_0_) to assess survival rates ^39^.

### Fluctuation test from a single-cell level of *E. coli* viability after UV irradiation (FT: fluctuation test)

To evaluate the contribution of primed cells to UV survival, we followed a method previously used for identifying primed cells under antibiotic treatment ^35^. An overnight *E. coli* culture was serially diluted (up to 10^10^-fold dilution) in a 96-well plate, with each well containing 200 μL. *E. coli* was cultured at 37°C until reaching log phase (OD_600_ of 0.3-0.5). In this approach, the 10^9^-10^10^ dilutions of *E. coli* overnight culture achieve single-cell level isolation. We selected 12 cultures, particularly from the 10^9^-10^10^ dilution conditions, for the UV inactivation experiment. These cultures were diluted 1000 times to prepare a 15 mL *E. coli* suspension (10^5^-10^6^ CFU/mL) in glass Petri dishes (Fig. 2). The UV irradiation conditions and the evaluation of UV-treated *E. coli* viability were identical to those described above.

### Evaluation of *E. coli* viability by sequential UV irradiation

*E. coli* suspension was prepared as described previously. First, the suspensions were treated with first UV irradiation, and then incubated in dark at room temperature for 30 min to allow for the induction and potential upregulation of DNA repair genes, including those regulated by the SOS response ^42^. Next, these suspensions were subjected to a second UV irradiation. Cell viability following sequential UV irradiation was assessed using the same method as described for the single-dose condition.

### Statement on the Use of Large Language Models (LLMs)

This manuscript was prepared with the assistance of a Large Language Model (LLM), specifically ChatGPT (OpenAI). All intellectual contributions and final approvals were made by the authors, who take full responsibility for the accuracy and integrity of the work.

## Results

### No detectable contribution of primed cells to survival under UV stress

The modified Luria–Delbrück fluctuation test was used to determine whether some bacterial cells are primed to withstand stress (Fig. 1). If tolerance is purely induced by UV irradiation, the coefficient of variation (CV) in viability between clonal populations in the fluctuation test (FT) and noise control (NC) will be similar. However, if certain cells are primed before UV irradiation, some clonal populations in FT will contain more primed cells than others, resulting in greater CV compared to NC (Fig. 1c).

A small fraction of stress-tolerant primed cells is present in the *E. coli* clonal population, and these cells exhibit high antibiotic tolerance ^35,36^. In this study, we aimed to evaluate whether primed cells contribute to survival under UV stress. We established an experimental framework for the modified fluctuation test in the UV inactivation of *E. coli*, following a previous study ^35^. Under FT condition, an overnight *E. coli* culture was serially diluted (up to 10^10^-fold) in a 96-well plate. At dilutions of 10^9^ to 10^10^, *E. coli* growth was observed in less than half of the wells (Fig. 2), indicating that single-cell level isolation was likely achieved ^35^. The log phase cultures in each well of the 96-well plate contained approximately 55 million *E. coli* cells on average, indicating that a single cell had undergone 25–26 generations of division. Each experiment involved testing 12 samples, and was replicated more than four times.

We applied a lower UV fluence of 2.5 mJ/cm², which serves as a permissive dose where UV-tolerant cells are more likely to appear ^39,43^. In the landmark study that identified primed cells, an antibiotic was applied under conditions that reduced the overall *E. coli* survival rate to ≤0.1%; nevertheless, some cultures prepared under FT conditions exhibited significantly higher viability ^35^. Drawing on this approach, we then employed a higher fluence of 6.0 mJ/cm², which reproducibly reduces *E. coli* viability to approximately 0.1% or less, to evaluate whether primed cells can confer protection under severe UV stress.

At a UV fluence of 2.5 mJ/cm^2^, *E. coli* viability ranged from 3.0 x 10^-3^ to 2.4 x 10^-1^ (Fig. 3a, Supplemental material 1). There was no significant difference in the viability or CV between NC and FT conditions (Figs. 3b and 3c). When the UV fluence was increased to 6.0 mJ/cm^2^, *E. coli* viability ranged from 5.0 x 10^-6^ to 6.1 x 10^-3^ (Fig. 3d, Supplemental material 2). There was no significant difference in the viability between NC and FT (Fig. 3e). CV in FT was significantly higher than in NC (Student’s *t*-test p = 0.040) (Fig. 3f). No significant correlation between the growth level (optical density) of *E. coli* culture and the *E. coli* viability after 6.0 mJ/cm^2^ UV irradiation was observed (Fig. 3g).

**Fig. 3.**
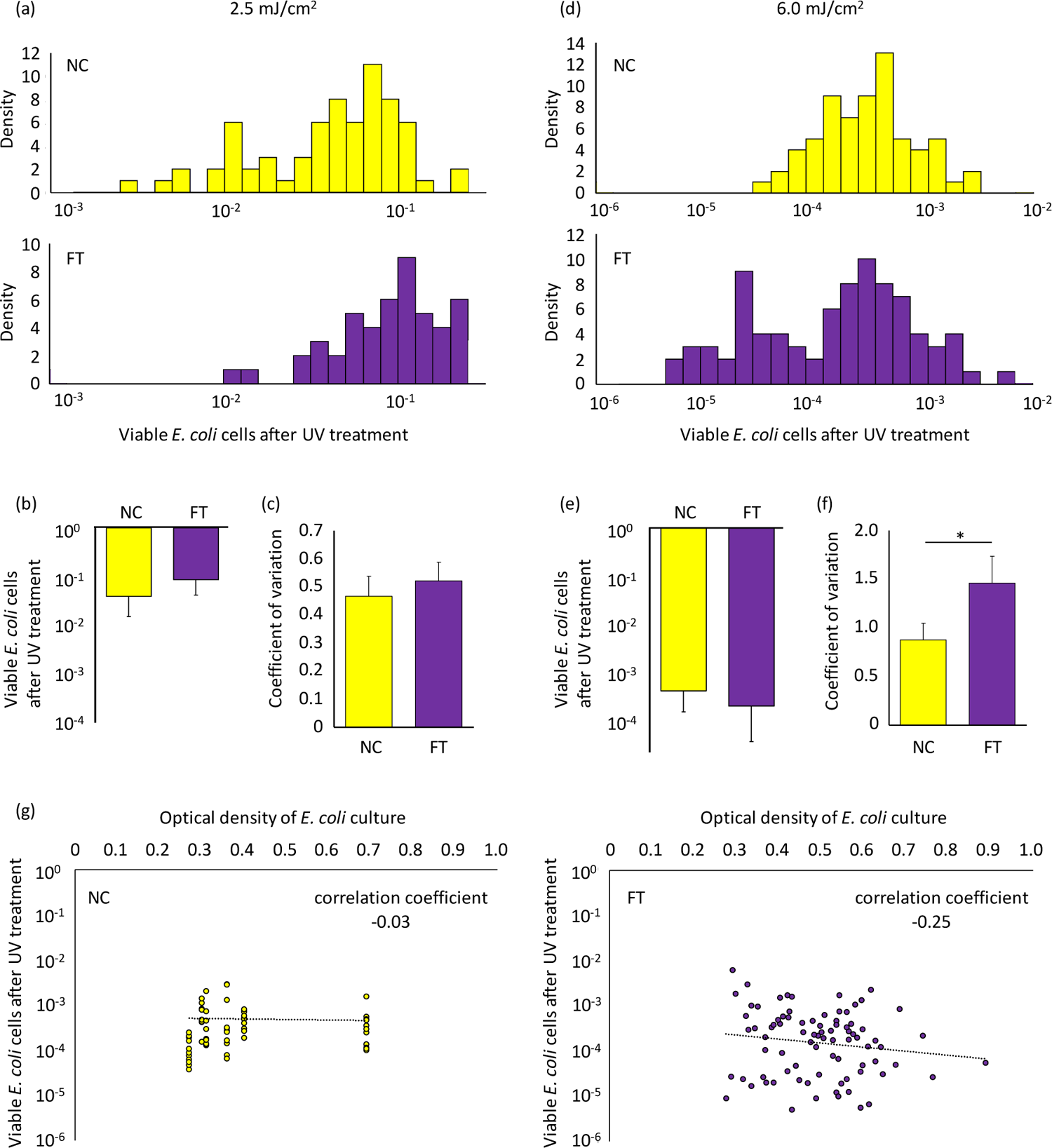
Variation of viable *E. coli* cells after the UV irradiation The *E. coli* suspensions were subjected to either 2.5 (a-c) or 6.0 (d-f) mJ/cm^2^ of 262 nm UV irradiation. (a and d) Histograms of viable *E. coli* cells after the UV irradiation. (b and e) Mean and (c and f) coefficient of variation of the viabilities observed in these experiments. Each experiment involved testing 12 technical replicates, and this was biologically replicated more than four times. Student’s *t*-test *p<0.05. Error bars indicate standard deviation. (g) The correlation between the growth level (optical density) of *E. coli* culture and the *E. coli* viability after 6.0 mJ/cm^2^ UV irradiation. Raw data are shown in Supplemental materials 1 and 2.

In the study by Hossain et al., the emergence of antibiotic-tolerant *E. coli* cells (primed cells) was concluded based on a significant increase in CV under FT conditions ^35^. Our investigation also revealed an elevated CV under FT conditions at a UV exposure of 6.0 mJ/cm^2^ (Fig. 3f), although this variance was not attributable to the appearance of highly UV-tolerant *E. coli* cells (Fig 3d).

To more rigorously assess the impact of primed cells on survival upon UV irradiation, we repeated the fluctuation test using a protocol in which every step—cultivation in mid-log phase, transfer to 96-well plates, and UV exposure—was identical for NC and FT conditions except for the initial cell number. This redesign approach minimized inoculum-related variability, thereby enabling a stricter evaluation of the priming hypothesis (Supplemental materials 3 and 4). Briefly, we grew both NC and FT in log phase in test tubes, and then transferred to 96 well plates (single cell for FT, and 10^4-5^ cells for NC) (Fig. S1). By the 6.0 mJ/cm^2^ UV irradiation, *E. coli* viability ranged from 2.9 x 10^-6^ to 3.7 x 10^-3^ (Fig. S2). The viability in NC was closely aligned with previous results (Fig. 3d and Fig. S2a). Although the overall viability distribution under FT conditions was similar to previous results, we observed a cluster of data around 10^-5^ to 10^-4^ (Fig. S2b). As a result, the *E. coli* viability in FT was significantly lower than that in NC (Student’s *t*-test p = 0.046) (Fig. S2c). We have previously reported the rare occurrence of UV-tolerant cells within *E. coli* clonal populations^39^. We speculate that these tolerant cells may enter a growth-arrested, tolerant mode. In FT conditions, each culture is derived from a single cell, potentially resulting in a relatively uniform growth pattern in the cell population. In the NC condition, cultures originate from 10^4-5^ cells. If cells prone to entering the tolerant mode exist within this population, it could lead to higher viabilities after the UV irradiation.

In our initial experiment, no subpopulation of highly UV-tolerant *E. coli* emerged under FT conditions (Fig. 3d). A follow-up experiment conducted within this study showed CV did not differ significantly between NC and FT cultures (Fig. S2d). Consequently, based on the data obtained, we could not confirm the existence of the primed cells identified by Hossain et al. that contribute to tolerance to UV stress in *E. coli*. Unlike the case of antibiotic tolerance ^35^, the concept of post-stress induction appears to better explain the emergence of surviving cells rather than pre-stress priming (Fig. 1).

### Reduced inactivation of *E. coli* by sequential UV irradiation at low fluence

UV irradiation causes DNA lesions in bacterial cells, impairing DNA replication and transcription, ultimately leading to cell death ^44^. The bactericidal effect of UV irradiation depends not only on the applied UV fluence but also on the bacterial capacity for DNA repair^45^. UV-induced DNA damage activates the SOS response ^46^, upregulating gene expressions involved in DNA repair systems, including nucleotide excision repair (NER) ^47^. In this study, we evaluated whether an initial UV exposure, which is expected to induce DNA repair systems, confers increased tolerance to subsequent UV irradiation.

The experimental conditions for UV exposure are illustrated in Fig. 4a. Upon UV irradiation at 2.5 and 3.5 mJ/cm^2^, the *E. coli* viability decreased to 5.6 × 10^-2^ and 2.0 × 10^-2^, respectively (Conditions 1 and 2 in Fig. 4b; Supplemental material 5). If cell survival were purely stochastic, the theoretical viability after sequential exposure to 2.5 and 3.5 mJ/cm^2^ would be expected to be 1.1 × 10^-3^, based on the multiplication of the individual survival fractions (Theoretical hypothesis in Fig. 4b). However, experimental results showed that the actual viability following sequential irradiation was 5.4 × 10^-3^ (Condition 3 in Fig. 4b), suggesting that the survival rate significantly higher than the predicted value (Student’s *t*-test p = 5.32 x 10^-6^).

**Fig. 4.**
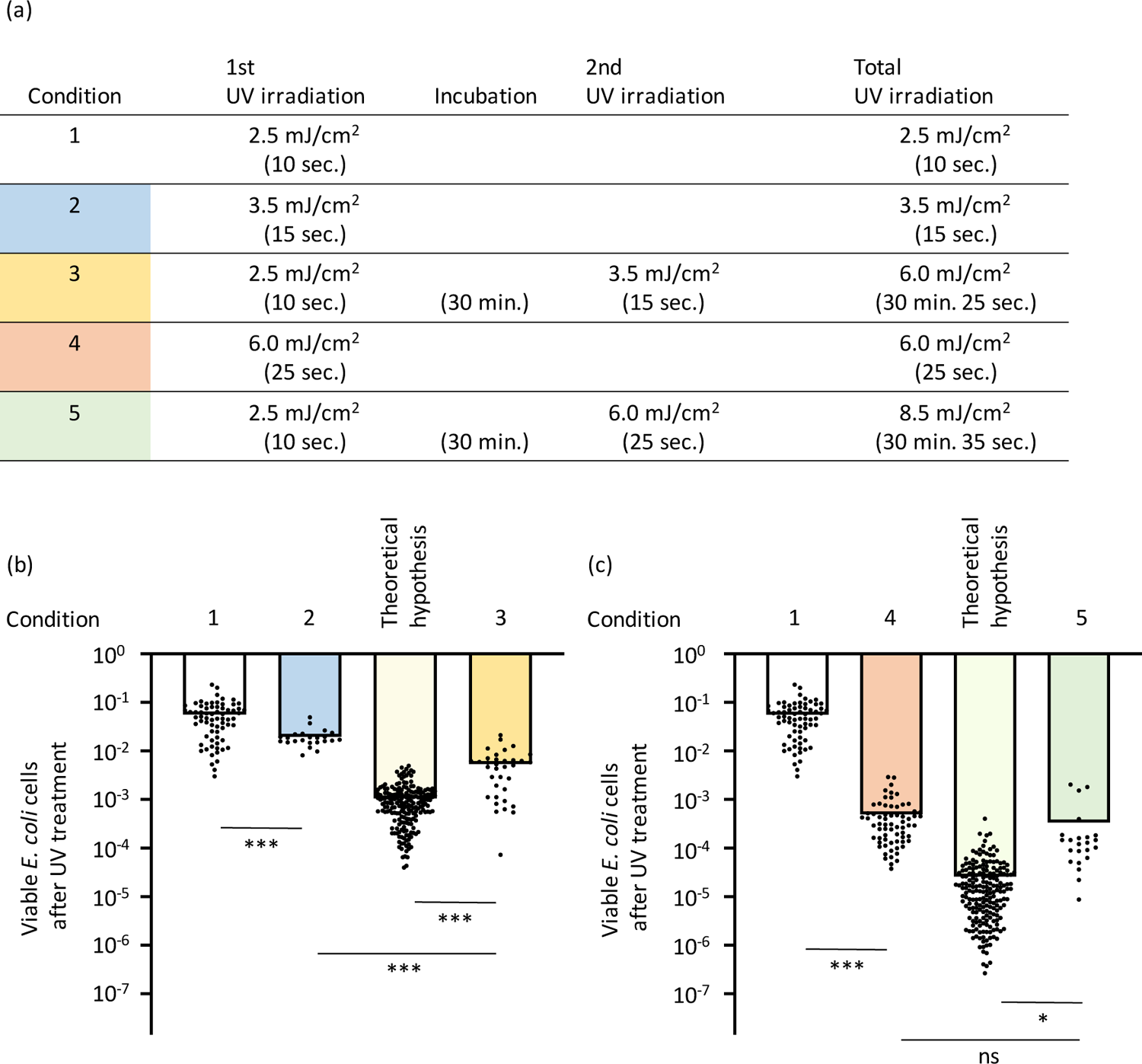
Effects of sequential UV irradiation on *E. coli* viability (a) Experimental setup for evaluating the effects of sequential UV irradiation. *E. coli* cells were exposed to UV fluences of 2.5, 3.5, and 6.0 mJ/cm^2^ using the 262 nm UV-LED device employed in this study. The exposure durations were 10, 15, and 25 seconds, respectively, as indicated in parentheses for each condition. (b) Viability of *E. coli* after UV irradiation under conditions 1, 2, and 3, as defined in panel (a). The expected theoretical viability was calculated as the product of the viabilities observed in conditions 1 and 2. This estimate was generated using a bootstrap approach based on the observed data from conditions 1 and 2. (c) Viability of *E. coli* after UV irradiation under conditions 3, 4, and 5, as defined in panel (a). The expected theoretical viability was calculated as the product of the viabilities observed in conditions 1 and 4, using the same bootstrap approach. Student’s *t*-test ***p<0.001. *p<0.05. Error bars indicate standard deviation. Raw data are shown in Supplemental material 5.

As mentioned above, when sequential irradiation (2.5 followed by 3.5 mJ/cm^2^) was applied, the total fluence amounted to 6.0 mJ/cm^2^, yielding a viability of 5.4 × 10^-3^ (Condition 3 in Fig. 4b). In contrast, single-dose irradiation with 6.0 mJ/cm^2^ resulted in a significantly lower viability of 5.0 × 10^-4^ (Condition 4 in Fig. 4c) (Student’s *t*-test p = 6.79 x 10^-7^). If survival were purely stochastic, the theoretical viability for sequential exposure to 2.5 and 6.0 mJ/cm^2^ would be 2.6 × 10^-5^ (Theoretical hypothesis in Fig. 4c). However, when 6.0 mJ/cm² of UV was applied following an initial 2.5 mJ/cm² exposure (Condition 5 in Fig. 4c), the observed viability was not significantly different from that of single-dose 6.0 mJ/cm² irradiation alone (Condition 4 in Fig. 4c) (Student’s *t*-test p = 0.274).

## Discussion

The concept of primed cells has been convincingly demonstrated under antibiotic treatment with ampicillin and apramycin ^35^. Ampicillin is a β-lactam antibiotic that disrupts bacterial cell wall synthesis by binding to and inactivating penicillin-binding proteins (PBPs)^48^. A fraction of the *E. coli* population can enter transient physiological states that upregulate stress responses, modify outer membrane permeability, or alter PBP expression ^49,50^. These changes may give rise to a “primed” phenotype, allowing these cells to survive lethal concentrations of ampicillin more effectively than the bulk population ^35^. Similarly, apramycin belongs to the aminoglycoside class, which targets the 30S ribosomal subunit to inhibit protein synthesis ^51^. Evidence suggests that *E. coli* can assume primed states by modulating efflux pumps, stress response regulators, and other components of the translational machinery ^52^. In both cases, these primed states are transient yet heritable for approximately seven generations, enabling short-term survival benefits without committing the entire population to a costly, constitutively tolerant phenotype ^35^.

This study indicated that such primed cells do not confer comparable benefits against UV irradiation. UV irradiation primarily damages DNA, creating lesions such as cyclobutane pyrimidine dimers and (6-4) photoproducts that disrupt replication and transcription ^44^. *E. coli* relies heavily on the SOS response to address such damage, coordinating the upregulation of DNA repair enzymes through the RecA–LexA regulatory circuit ^53^. The expression of genes involved in the SOS response is strongly induced upon DNA damage caused by UV irradiation ^54^.

If a subpopulation were primed for UV stress in a manner similar to antibiotic tolerance, we would expect them to preemptively activate key SOS genes or DNA repair enzymes prior to UV irradiation. However, our fluctuation test did not detect a distinct subpopulation with significantly higher survival after UV treatment, even under conditions designed to reveal rare, highly tolerant phenotypes (Fig. 3). It is important to note, however, that this assay may not be able to capture primed states with very short-term memory. If the transient memory lasts only one to two generations, it may result in no detectable difference between FT and NC. Taking this into account, the fluctuation test suggests an absence of long-term cellular priming—such as the approximately seven-generation memory reported for antibiotic tolerance—but does not rule out the possibility of rapidly decaying, short-term primed states prior to UV stress (Fig. 1).

Antibiotics typically exert stress by targeting specific cellular structures or processes, allowing for targeted molecular adaptations (e.g., changes in membrane permeability or ribosomal proteins) ^50,51^. UV radiation causes widespread DNA lesions, and successful tolerance often requires robust DNA repair mechanisms that are activated immediately after damage ^44^. If repair enzymes are not already upregulated, even a slight delay can result in lethal double-strand breaks or replication fork collapse ^55^. Thus, *E. coli* may not sustain a stable “primed” state for UV stress as partial induction of DNA repair processes is likely insufficient to handle the sudden, substantial DNA insult inflicted by UV irradiation. Our observations support the idea that tolerance to antibiotics and UV stress may operate via partially distinct mechanisms that merit separate investigation.

The experimental results of sequential UV irradiation showed higher bacterial viabilities than predicted by the theoretical stochastic model, which was calculated by multiplying individual survival fractions (Fig. 4). The concept of post-stress induction better explains the emergence of tolerance to UV stress (Figs. 1 and 3). In our previously repeated UV treatment experiments, the surviving cells—after regrowth in fresh medium—did not consistently show higher tolerance in subsequent exposures ^39^, suggesting that the observed tolerance is a transient, induced response rather than the results of genetic resistance ^40,41^. During sequential UV exposures, the induction of DNA repair systems may mitigate cumulative inactivation more effectively than a single-dose exposure with an equivalent total fluence. During sequential irradiation, each partial dose may remain within the threshold that bacterial repair systems can effectively counteract, thereby reducing the overall inactivation rate. Bacterial inactivation curves under UV irradiation often exhibit an initial “shoulder” phase, characterized by delayed inactivation, followed by a log-linear decline ^43^. This shoulder effect is widely attributed to DNA repair processes that mitigate damage at low UV doses. *E. coli* strains with proficient DNA repair capabilities exhibit pronounced shoulders in their UV inactivation curves, whereas repair-deficient mutants show a more linear inactivation pattern^56^. These findings suggest that sublethal DNA lesions can be repaired by mechanisms such as nucleotide excision repair (NER), allowing cells to survive until the repair capacity is exceeded. Once the threshold for maximum DNA repair is surpassed, additional UV exposure results in rapid exponential cell death ^43^. Thus, the shoulder effect primarily reflects active DNA repair processes that buffer the impact of UV exposure.

In future investigations, it would be valuable to confirm and refine our conclusion that survival under UV stress arises solely through induced mechanisms rather than any primed state. Direct measurement of the induction levels of SOS genes—such as *recA* and *uvrA*— through quantitative PCR or transcriptomic profiling would provide molecular validation for the proposed induced response model. More detailed single-cell lineage tracking using time-lapse fluorescence microscopy could help visualize the real-time expression of SOS genes in individual cells ^39,57,58^. This approach could definitively clarify whether a transient, epigenetically heritable state is ever induced prior to UV exposure—or if, indeed, such priming does not occur under any tested conditions. In parallel, emerging single-cell omics technologies offer the potential to dissect transcriptional and epigenetic modifications in unprecedented detail, identifying specific markers that might either confirm the absence of primed cells or reveal overlooked transient subpopulations ^12,59,60^. By integrating these advanced methodologies, future research can more conclusively define the limits of bacterial adaptation to UV stress and help develop new strategies to mitigate the impact of stress-induced survival.

## Acknowledgements

We sincerely thank Prof. Hideto Miyake, Prof. Norihiro Nishimura, and Dr. Mina Okamura of Mie University for their cooperation in providing access to the UV-LED device. Institute for Fermentation, Osaka provided partial funding for this study. This work is also supported by NIH-NIGMS grant R35GM148351. These organizations had no involvement in designing the research, collecting or interpreting data, or deciding to publish the results.

## Author contributions

S.I., Y.S., and A.S. conceptualized and designed the study. S.I., M.T., J.T., and E.M. carried out the data collection, analysis, and interpretation. S.I. and A.S. prepared and revised the manuscript. All authors reviewed and approved the final version for submission.

## Abbreviations

LB: Luria-Bertani
CFU: colony-forming unit
UV-LED: ultraviolet light-emitting diode
NC: noise control
FT: fluctuation test
CV: coefficient of variation
NER: nucleotide excision repair

