## Supplemental material 3 for "Induced tolerance to UV stress drives survival heterogeneity in isogenic *E. coli* cell populations"

**MATERIALS AND METHODS**

*Escherichia coli* K-12 freeze stock was used as the inoculum and aerobically cultured in 5 mL of Luria-Bertani (LB) broth at 37 °C and 220 rpm in a test tube for 12 hours. The culture was then inoculated into fresh 5 mL LB broth and aerobically cultured under the same conditions until reaching the log phase (OD_660_ of 0.3-0.5), corresponding to 10^8^-10^9^ *E. coli* cells/mL.

The log phase culture was diluted with 200 μL of fresh LB broth in each well of a 96-well plate. For the noise control (NC) condition, the culture was diluted 10^4^-fold, resulting in approximately 10^4^-10^5^ *E. coli* cells per well. For the fluctuation test (FT) condition, the culture was diluted 10^8^-10^9^-fold to achieve one *E. coli* cell per well (Fig. S1).

Cell growth was monitored using a Stratus plate reader (Cerillo, VA, USA) at 37 °C and 150 rpm. The log phase culture (OD_600_ of 0.2-0.3) was diluted 1000-fold in phosphate-buffered saline to prepare 15 mL suspensions of *E. coli* (10^5^-10^6^ CFU/mL) in 55 mm diameter glass Petri dishes. Eight to twelve samples were prepared for each experiment. A UV-LED (262 nm, Nikkiso Giken Co. Ltd., Ishikawa, Japan) was positioned at 32 mm above the surface of the *E. coli* suspension. The *E. coli* suspensions were subjected to 6.0 mJ/cm² of 262 nm UV irradiation with 700 rpm stirring. After the UV irradiation, the UV-treated *E. coli* was cultured on 3 M Petrifilm Aerobic Count Plate (3 M Company, MN, USA) at 35 °C for 48 h. We counted viable *E. coli* cells as colony-forming units (CFUs) from a 1 mL suspension. Viable cell counts (CFU/mL) in UV-treated samples (N) were normalized against those in untreated samples (N_0_).

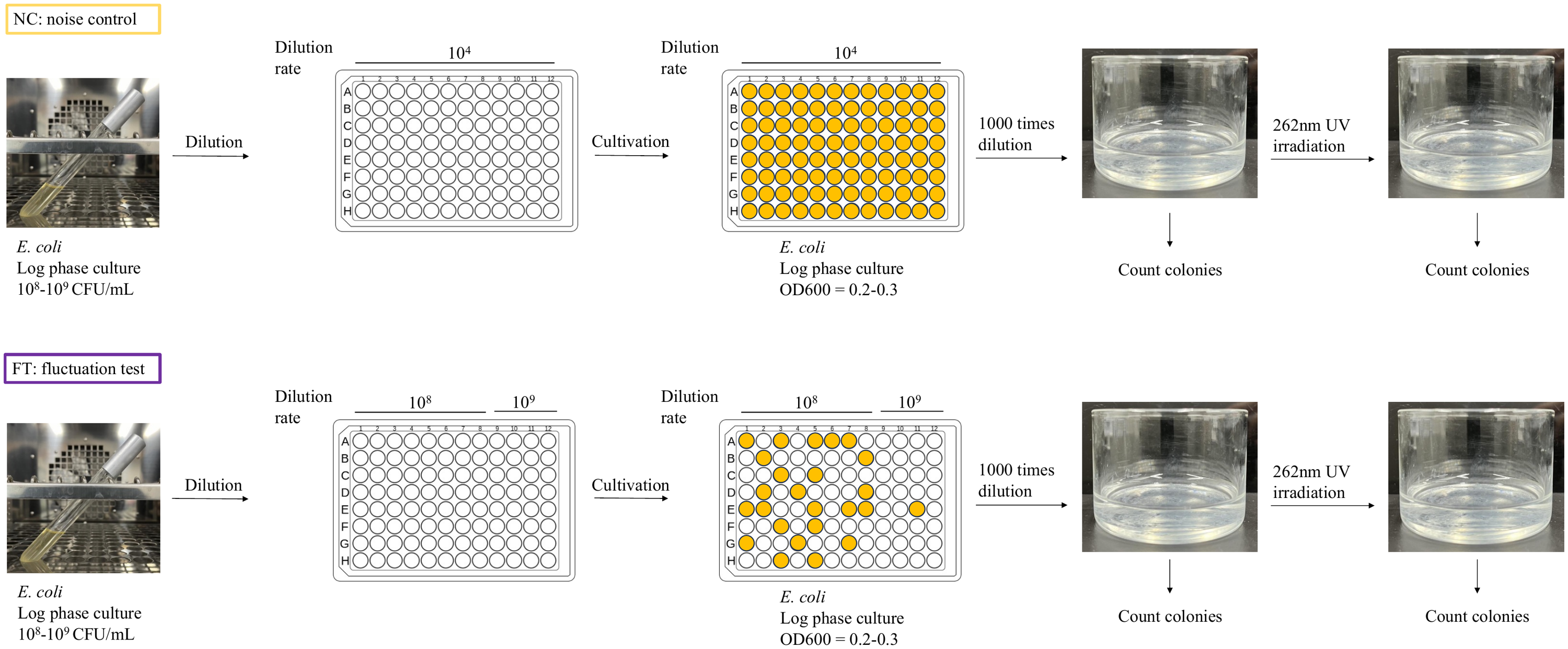

Fig. S1: Experimental design for the fluctuation test by UV irradiation

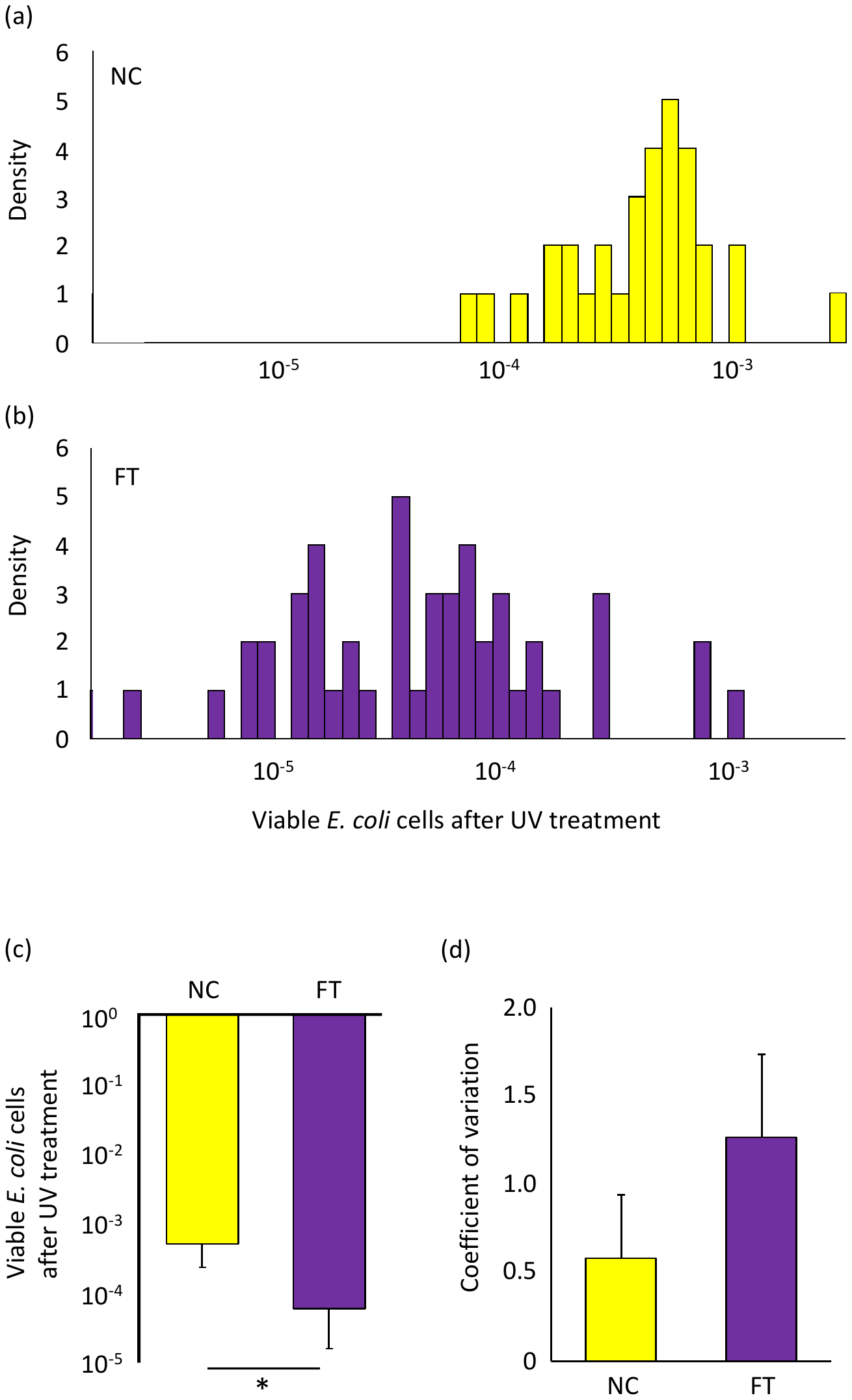

Fig. S2: Viability of *E. coli* cells after the UV irradiation

(a and b) Histograms of viable *E. coli* cells after the UV irradiation. (c) Mean and (d) coefficients of variance observed in these experiments. Error bars indicate standard deviations. *p<0.05.
